# Comparison of multiple transcriptomes exposes unified and divergent features of quiescent and activated skeletal muscle stem cells

**DOI:** 10.1101/219998

**Authors:** Natalia Pietrosemoli, Sébastien Mella, Siham Yennek, Meryem B. Baghdadi, Hiroshi Sakai, Ramkumar Sambasivan, Francesca Pala, Daniela Di Girolamo, Shahragim Tajbakhsh

## Abstract

**Background:** Skeletal muscle stem cells (MuSCs) are quiescent in adult mice and can undergo multiple rounds of proliferation and self-renewal following muscle injury. Several labs have profiled transcripts of myogenic cells during developmental and adult myogenesis with the aim of identifying quiescent markers. Here, we focused on the quiescent cell state and generated new transcriptome profiles that include subfractionations of adult MuSC populations, and an artificially induced prenatal quiescent state, to identify core signatures for quiescent and proliferating MuSCs.

**Methods:** Comparison of available data offered challenges related to the inherent diversity of datasets and biological conditions. We developed a standardized workflow to homogenize the normalization, filtering, quality control steps for the analysis of gene expression profiles allowing the identification up- and down-regulated genes and the subsequent gene set enrichment analysis. To share the analytical pipeline of this work, we developed Sherpa, an interactive Shiny server that allows multiscale comparisons for extraction of desired gene sets from the analysed datasets. This tool is adaptable to cell populations in other contexts and tissues.

**Results:** A multiscale analysis comprising eight datasets of quiescent MuSCs had 207 and 542 genes commonly up- and down-regulated, respectively. Shared up-regulated gene sets include an over-representation of the TNFa pathway via NFKb signaling, Il6-Jak-Stat3 signaling, and the apical surface processes, while shared down-regulated gene sets exhibited an over-representation of *Myc* and *E2F* targets, and genes associated to the G2M checkpoint and oxidative phosphorylation. However, virtually all datasets contained genes that are associated with activation or cell cycle entry, such as the immediate early stress response genes *Fos* and *Jun*. Empirical examination of fixed and isolated MuSCs showed that these and other genes were absent *in vivo*, but activated during procedural isolation of cells.

**Conclusions:** Through the systematic comparison and individual analysis of diverse transcriptomic profiles, we identified genes that were consistently differentially expressed among the different datasets and common underlying biological processes key to the quiescent cell state. Our findings provide impetus to define and distinguish transcripts associated with true *in vivo* quiescence from those that are first responding genes due to disruption of the stem cell niche.

## Background

Most adult stem cell populations identified to date are in a quiescent state [1]. Following tissue damage or disruption of the stem cell niche, skeletal muscle stem (satellite) cells (MuSCs) transit through different cell states from reversible cell cycle exit to a postmitotic multinucleate state in myofibres. In mouse skeletal muscle, the transcription factor *Pax7* marks MuSCs during quiescence and proliferation, and it has been used to identify and isolate myogenic populations from skeletal muscle [2, 3]. Myogenic cells have also been isolated by Fluorescence Activated Cell Sorting (FACS) using a variety of surface markers, including α7-integrin, VCAM and CD34 [4] Although these cells have been extensively studied by transcriptome, and to a more limited extend by proteome profiling, different methods have been used to isolate and profile myogenic cells thereby making comparisons laborious and challenging. To address this issue, it is necessary to generate comprehensive catalogs of gene expression data of myogenic cells across distinct states and in different conditions.

Soon after their introduction two decades ago, high-throughput microarray studies started to be compiled into common repositories that provide to the community access to the data. Several gene expression repositories for specific diseases, such as the Cancer Genome Atlas (TCGA) [5], the Parkinson’s disease expression database ParkDB [6], or for specific tissues, such the Allen Human and Mouse Brain Atlases [7][8] among many, have been crucial in allowing scientists the comparison of datasets, the application of novel methods to existing datasets, and thus a more global view of these biological systems.

In this work, we generated transcriptome datasets of MuSCs in different conditions and aimed to perform comparisons with published datasets. Due to the diversity of platforms and formats of published datasets, this was not readily achievable. For this reason, we developed an interactive tool called *Sherpa* (SHiny ExploRation tool for transcriPtomic Analysis) to provide comprehensive access to the individual datasets analysed in a homogeneous manner. This webserver allows users to: i) identify differentially expressed genes of the individual datasets, ii) identify the enriched gene sets of the individual datasets, and iii) effectively compare the chosen datasets. *Sherpa* is adaptable and serves as a repository for the integration and analysis of future transcriptomic data. It has a generic design that makes it applicable to the analysis of other transcriptome datasets generated in a variety of conditions and tissues.

We analyse gene expression profiles (GEPs) of activated and quiescent states of mouse MuSCs derived from three new experimental setups and six publicly available microarray datasets to define a consensus molecular signature of the quiescent state. This large compendium of expression data offers the first comparison and integration of nine independent studies of the quiescent state of mouse satellite cells and we developed Sherpa, a shiny interactive web server to provide a user-friendly exploration of the analysis. In addition, using a protocol for the fixation and capture of mRNA directly from the tissue without the alteration in gene expression that could arise during the isolation procedure, which typically takes several hours with solid tissues, we have empirically tested the expression of transcripts. Strikingly, several genes, including members of the *Jun* and *Fos* family were found to be present in isolated MuSCs using conventional isolation procedures, but they were absent *in vivo*. These findings, and the unique atlas that we report, will undoubtedly improve our current understanding of the molecular mechanisms governing the quiescent state and contribute to the identification of critical regulatory genes involved in different cell states.

## Methods

### Individual dataset transcriptomic analysis

The analysis comprised a total of nine datasets, three novel microarray datasets and six publicly available datasets [9][10][11][12][13][14], choosing only samples with overall similar conditions. All datasets were analysed independently following the same generalised pipeline based on ad-hoc R implemented scripts (Fig. 2).

### Gene expression profiles

The microarray data compared activated satellite cells (ASCs) and quiescent satellite cells (QSCs) from different experiments. Table 1 describes the public datasets that were taken into account for the analysis with the GEO [15] (Gene Expression Omnibus) identifications, references and sample distribution. The new mouse microarray datasets include the following comparisons: young adult Quiescent(adult) / Activated (postnatal day 8), and Quiescent [high/low] / D3Activated [high/low], and Foetal_NICD [E17.5/E14.5]. Table 1 presents the sample details.

**Table 1.** Summary of analysed transcriptomic datasets of activated and quiescent states of mouse muscle stem (satellite) cells. Three high-throughput experimental setups and six publically available microarray datasets comparing activated satellite cells (ASCs) and quiescent satellite cells (QSCs) are shown in the rows. The biological, experimental and technical details of each experiment are shown in the different columns of the Table. (h=hours, d=days, w=weeks, m=months).

| Ref/Code | Quiescent [high/low] 3D Activated [high/low] | Quiescent Activated | Foetal R26<sup>NICD</sup> [E17.5/E14.5] | GSE47177 Liu et al [9] | GSE3483 Fukada et al [10] | GSE15155 Pallafacchina et al [11] | GSE38870 Farina et al. [12] | GSE70376 García-Prat et al. [13] | GSE81096 Lukjanenko et al. [14] |
| --- | --- | --- | --- | --- | --- | --- | --- | --- | --- |
| Num. of samples | 3 QSC_Pax 7 low, 3 ASC_Pax 7 low, 3 QSC_Pax 7 high, 3 ASC_Pax 7 high | 3 QSC, 3 P8_ASC | 3 eq_QSC, 3 eq_ASC | 3 QSC, 3 ASC 60h, 3 ASC 84h | 3 QSC, 3 ASC | 3 QSC, 3 ASC | 3 QSC, 3 ASC | 4 QSC, 4 ASC | 6 QSC, 5 ASC |
| Date | 2013 | 2007 | 2015 | 2013 | 2007 | 2010 | 2012 | 2015 | 2016 |
| Anatomy | Tibialis anterior | Limb, bodywall, diaphragm | Forelimbs | Hindlimb | Hindlimb | Diaphragm, pectorales, abdominal muscles | Tibialis anterior | Tibialis anterior | Tibialis anterior, gastrocnemius, quadriceps |
| Sex | M |  | M,F | M | F | M,F | F | M | M |
| Age | 6-8 w | P8, 4-5 w | E14.5, E17.5 | 8 w | 8-12 w | 6 w | 3-6 m | Young: 3 m, old: 20-24 m | Young: 9-15 w, old: 20-24 m |
| Strain | C57BL/6 | B6.129 |  | C57BL/6 (Jackson) | C57BL/7 (nihon clea) |  | C57BL/6 x DBA2 (??) | C57BL/6 | C57BL/6 (Janvier) |
| Reporter | Tg:Pax7-nGFP (10% high, 10% low) | Pax7<sup>nGFP/+</sup> | Myf5<sup>Cre+</sup>: R26<sup>stop-NICDgfp/+</sup> | Pax7<sup>CreER/+</sup>: R26<sup>eYFP/+</sup> |  | Pax3<sup>GFP/+</sup> (high GFP) |  |  |  |
| Activation | Notexin | P8 = "Activated" | E14.5 = "Activated" | BaCl<sub>2</sub> | QSCs in culture for 4 d | Pax3<sup>GFP/+</sup>, Pax3<sup>GFP/+</sup>:mdx:mdx<br>Adult; Adult mdx; 1 w old; 3 d in culture | Injury: BaCl<sub>2</sub> (50μL 1.2%) | Cardiotoxin | Cardiotoxin |
| QSCs Purif. | Reporter | Reporter | Reporter | FACS: Pax7<sup>CreER/+</sup>: R26<sup>eYFP/+</sup> | FACS: CD45-/SM/C-2.6+ | Reporter | FACS: Syndecan-3 | FACS: integrin-alpha7+ /Lin-/CD31-/CD45-/CD11b-/Sca1- | CD34+/ integrin-alpha7+/Lin- |
| Timing | Quiescent & 3 d post-injury |  | See age | 36h (1.5d), 60h (2.5 d), 84h (3.5d) post-injury | 4 d in culture | 3 d in culture | 12h or 48h post-injury | 72h (3d) | 72h (3d) |
| Platform | Affymetrix Mouse Gene 1.0ST | Affymetrix 430_2.0 | Affymetrix Mouse Gene 1.0ST, Affymetrix Mouse Gene 2.0ST | Affymetrix Mouse Gene 1.0ST | Affymetrix 430A | Affymetrix 430_2.0 | Affymetrix 430_2.0 | Agilent 028005 SurePrint G3Mouse 8x60k Microarray | Illumina MouseRef-8 v2.0 |

### Animals, injuries and cell sorting

Animals were handled according to national and European Community guidelines, and an ethics committee of the Institut Pasteur (CTEA) in France. For isolation of quiescent MuSCs, *Tg: Pax7-nGFP* mice (6-12 weeks) [2] were anesthetized prior to injury. *Tibialis antorior* (TA) muscles were injured with notexin (10μl – 10μM; Latoxan). Cells were then isolated by FACS using BD FACS ARIA III, MoFlo Astrios and Legacy sorters. Pax7^Hi^ and Pax7^Lo^ cells correspond to the 10% of cells with the highest and the lowest expression of nGFP, respectively, as defined previously [3].

For isolation of activated MuSCs, TA muscles (day 3 post-injury (D3) and non-injured) were collected and subjected to 4-5 rounds of digestion in a solution of 0.08% Collagenase D (Roche) and 0.1% Trypsin (Invitrogen) diluted in DMEM-1% P/S (Invitrogen) supplemented with DNAse I at 10μg/ml (Roche) [2][3]. Pax7^Hi^ and Pax7^Lo^ cells correspond to the 10% of cells with the highest and the lowest expression of nGFP, respectively, as defined previously [3].

Skeletal muscle progenitors were obtained also from the forelimbs of E14.5 and E17.5 foetuses of *Myf5 ^CreCAP/+^:R26R ^stop-NICD-nGFP/+^* [16] compound mice. Tissues were dissociated in DMEM (GIBCO, 31966), 0.1% Collagenase D (Roche, 1088866), 0.25% trypsin (GIBCO, 15090-046), DNase 10 μg/ml (Roche, 11284932001) for three consecutive cycles of 15 min at 37°C in a water bath under gentle agitation. For each round, supernatant containing dissociated cells was filtered through 70μm cell strainer and trypsin was inhibited with calf serum. Pooled supernatants from each round of digestion were centrifuged at 1600rpm for 15 min at 4°C and pellet was re-suspended in cold DMEM/1% PS/2%FBS and filtered through 40μm cell strainer.

In other experiments, skeletal muscles from the limbs, body wall and diaphragm were collected from pups at postnatal day 8 (P8, mitotically active satellite cells) and 4-5 weeks old mice (quiescent satellite cells) of *Pax7 ^nGFP/+^* knock-in line [17]. Cells were isolated by FACS based on NICD-GFP or Pax7-nGFP intensity, using BD FACS ARIA III (BD Biosciences) and MoFlo Astrios (Beckman Coulter) sorters.

### Microarray sample preparation

Total mRNAs were isolated using (Qiagen RNAeasy^®^ Micro Kit) according to the manufacturer’s recommendations; 5 ng of total RNA was reverse transcribed and amplified following the manufacturer's protocols (Ovation Pico WTA System v2 (Nugen Technologies, Inc. #3302-12); Applause WTA Amp-Plus System (Nugen Technologies, Inc. #5510-24)), fragmented and biotin labelled using the Encore Biotin Module (Nugen Technologies, Inc. #4200-12). Gene expression was determined by hybridization of the labelled template to Genechip microarrays Mouse Gene 1.0 ST (Affymetrix). Hybridization cocktail and post-hybridization processing was performed according to the “Target Preparation for Affymetrix GeneChip Eukaryotic Array Analysis” protocol found in the appendix of the Nugen protocol of the fragmentation kit. Arrays were hybridized for 18 hours and washed using fluidics protocol FS450 0007 on a GeneChip Fluidic Station 450 (Affymetrix) and scanned with an Affymetrix Genechip Scanner 3000, generating CEL files for each array. Three biological replicates were run for each condition.

### Western blot analysis

Total protein extracts from satellite cells isolated by FACS were run on a 4-12% Bis-Tris Gel NuPAGE (Invitrogen) and transferred on Amersham Hybond-P transfer membrane (Ge Healthcare). The membrane was then blocked with 5% nonfat dry milk in TBS, probed with anti-JunD (329) (1:1000, sc-74 Santa Cruz Biotechnology Inc.), anti-JunB (N-17) (1:1000, sc-46 Santa Cruz Biotechnology Inc.) or anti-c-Jun (H-79) (1:1000, sc-1694 Santa Cruz Biotechnology Inc.) overnight, washed and incubated with HRP-conjugated donkey anti-rabbit IgG secondary antibody (1:3000), and detected by chemiluminescence (Pierce ECL2 western blotting substrate, Thermo Scientific) using the Typhoon imaging system. After extensive washing, the membrane was incubated with anti-Histone H3 antibody (ab1691, 1:10000; abcam) as loading control. All Western blots were run in triplicate and bands were quantitated in 1 representative gel. Quantification was done using ImageJ software.

### Isolation of fixed mouse muscle stem cells and real-time PCR

For empirical analysis of genes by RT-qPCR (e.g. *Jun* and *Fos*), skeletal muscles were fixed immediately in 0.5% for 1 h in paraformaldehyde (PFA) using a protocol based on the notion that transcripts are stabilized by PFA fixation [18] [Machado et al., in press; P. Mourikis and F. Relaix, personal communication]. Briefly, PFA fixed and unfixed skeletal muscles were minced as described [4], fixed samples were incubated with collagenase at double the normal concentration and mRNA was isolated following FACS based on size, granulosity and GFP levels using a FACS Aria II (BD Bioscience). Total RNA was extracted from fixed cells with RecoverAll™ Total Nucleic Acid Isolation Kit Ambion, ThermoFisher), according to manufacturer instructions. cDNA was prepared by random-primed reverse transcription (Super-Script II, Invitrogen, 18064-014), and real-time PCR was done using SYBR Green Universal Mix (Roche, 13608700) StepOne-Plus, Perkin-Elmer (Applied Biosystems). Specific primers for each gene were designed, using the Primer3Plus online software, to work under the same cycling conditions. For each reaction, standard curves for reference genes were constructed based on six 4-fold serial dilutions of cDNA. All samples were run in triplicate. The relative amounts of gene expression were calculated with RLP13 expression as an internal standard (calibrator). RT-qPCR primers used appear in Table S2.

### Normalisation, quality control and filtering of GEPs

GEPs were processed using standard quality control tools to obtain normalised, probeset-level expression data. For all raw datasets derived from affymetrix chips, Robust Multi-Array Average expression measure (rma) was used as normalization method using the *affy* and the *oligo* R packages [19][20]. All analyses were preferentially conducted at the probeset level. Probesets were annotated to gene symbol and gene ENTREZ using chip-specific annotations. For gene level results, the probeset with the highest expression variability was selected to represent the corresponding gene. Quality controls were performed on raw data using Relative Log Expression (RLE) and Normalised Unscaled Standard Errors (NUSE) plots from the *affyPLM* R package [21]. Sample distribution was examined using hierarchical clustering of the Euclidean distance and Principal Component Analysis from the *stats* [22] and *FactoMineR* R packages [23] (See Additional file 1: Fig. S1 for the resulting plots for dataset Quiescent [high/low] / D3Activated [high/low]). The resulting plots of the remaining datasets are not shown but they show similar trends, which can be explored through the interactive webserver Sherpa.

### Differential expression analysis

Each dataset was individually analysed to identify genes showing significant differential expression (DEGs) between the ASC and the QSC (Gene level analysis in Fig. 1; Differential analysis in Fig. 2). This analysis was performed using the linear model method implemented in the *Limma* R package [24]. The basic statistic was the moderated t-statistic with a Benjamini and Hochberg’s multiple testing correction to control the false discovery rate (FDR) [25].

**Fig. 1.**
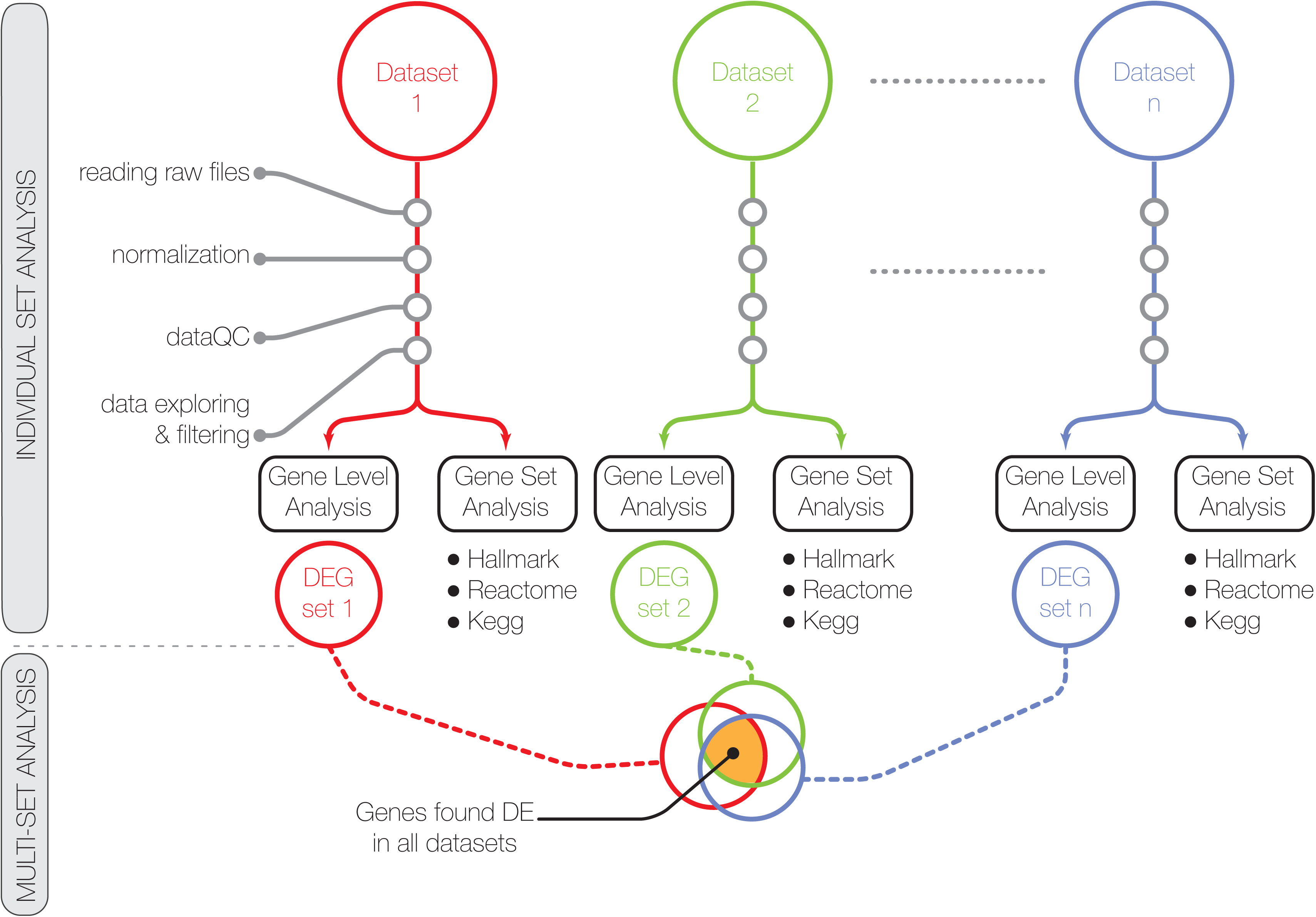
General framework of the analysis: an individual dataset analysis followed by a multi-set analysis. The individual dataset analysis consisted of: i) the analysis of gene expression profiles (GEPs) of each dataset, including normalisation, filtering and quality control check of each raw dataset, and the differential analysis to identify dataset-specific differentially expressed genes (DEGs); ii) the Gene set enrichment analysis (GSEA) performed in the gene set space. The GSEA consisted in identifying enriched pathways from three gene sets of the MSigDB collection [27] (Hallmark gene sets, CP:KEGG gene sets and CP: Reactome gene sets); iii) a multi-set analysis to assemble a study-independent gene signature, i.e. a list of genes specific to the quiescence state.

### Gene set enrichment analysis on individual sets

Each dataset was tested for gene set enrichment independently, using the CAMERA competitive test implemented in the Limma R package [30], and three gene set collections from the mouse version of the Molecular Signatures Database MSigDB v6 [26][27]: 1) Hallmark gene sets (H), which summarize and represent specific well-defined biological states or processes displaying a coordinate gene expression, 2) KEGG canonical pathways (C2 CP:KEGG), derived from the Kyoto Encyclopedia of Genes and Genomes [28] and 3) Reactome canonical pathways (C2 CP:Reactome) from the curated and peer reviewed pathway database [29] (Gene set analysis in Figs. 1 and 2).

**Fig. 2.**
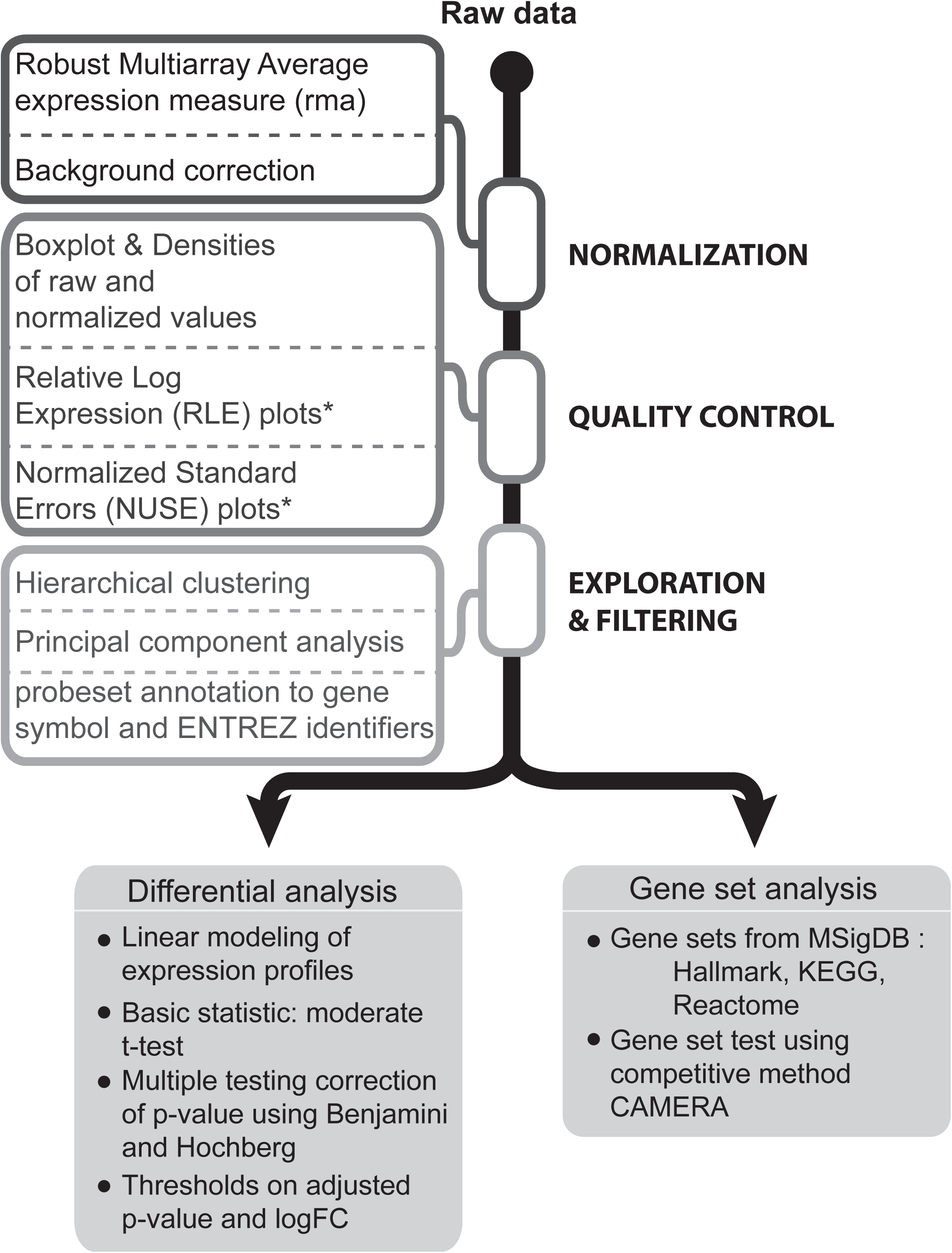
Workflow of the standardized individual dataset analysis. The analysis of the nine datasets was performed in a consistent manner for each dataset using ad-hoc R scripts. It included a first step of data preparation followed by a second step of data analysis. GEPs were processed using standard quality control tools to obtain normalised, probeset-level expression data. For raw datasets derived from affymetrix chips, Robust Multi-Array Average expression measure (rma) was used as normalization method. All analyses were conducted at probeset level. Probesets were annotated to gene symbol and gene ENTREZ using chip-specific annotations. Quality controls were performed on raw data using RLE and NUSE plots. The distribution of the QSC and ASC samples according to their GEPs was explored using hierarchical clustering of the Euclidean distance and Principal Component Analysis (Additional file 1: Figure S1). Statistically differentially expressed genes (DEGs) were identified between the ASC and the QSC groups using the linear model implemented by the Limma R package [10]. Gene set enrichment analysis was based on three gene set collections from the mouse version of the Molecular Signatures Database MSigDB v6.0 [12][13]: 1) Hallmark, which summarizes and represents specific well-defined biological states or processes displaying a coordinate gene expression, 2) KEGG canonical pathways, derived from the Kyoto Encyclopedia of Genes and Genomes [14] and 3) Reactome canonical pathways from the curated and peer reviewed pathway database [15]. To test for the enrichment of these gene sets, the competitive gene set test CAMERA [16] was used.

### Multiple set analysis: determination of the quiescent signature

The combinatorial landscape of datasets was explored using the *SuperExactTes*t [31] and the *UpSetR* [32] R packages to test and visualize the intersection of the datasets. Additionally, the Jaccard index [33] of similarity was calculated to assess the extent of similarity between DEGs of each pair of datasets. A significance ranking, based on several criteria, was calculated for each individual dataset to determine its presence or absence in the final dataset ensemble, which was used for determining the gene signature. Once the dataset ensemble was defined, the overlapping differentially up- and down-regulated genes (DEGs, as defined by the adjusted p-value ≤ 0.05) were used to build the quiescent signature.

### Gene set enrichment analysis on the quiescent signature

An over-representation analysis (ORA) [34] was applied to the quiescent signature using the previously described gene collection (Hallmark, Kegg, Reactome). For this purpose, commonly up-regulated or down-regulated genes were used in a one-sided Fisher’s exact test implemented in R script with a Benjamini and Hochberg’s multiple testing correction of the p-value to determine the enriched gene sets and the direction of such enrichment.

### Web application: Sherpa

We developed an interactive web application for the exploration, analysis and visualization of the individual datasets and their combination (http://sherpa.pasteur.fr). This application allows the user to effectively and efficiently analyse the individual datasets one by one (individual dataset analysis) or as an ensemble of datasets (multi-set analysis) and was developed with the *Shiny* R package [35].

## Results

This study involves an individual dataset analysis followed by a multi-set analysis (Fig. 1). First, each raw dataset was normalised, filtered and subjected to the same quality controls and checks. Gene level differential analysis and gene set enrichment analysis were then performed (Fig. 2). Finally, a multi-set analysis assembled a platform-independent list of genes specific to the quiescent state. When analysing multiple microarray GEPs, however, several issues needed to be addressed regarding the experimental set-up, the microarray platforms and the laboratory conditions [36]. First, the individual studies, even if related, had different aims, experimental designs and cell populations of interests (e.g. developmental stage, and gender of mice). Second, the different microarray platforms contained different probes and probesets with specific locations and alternative splicing that might produce different expression results [37]. Finally, sample preparation, protocols and dates of extractions might have influenced array hybridization and introduced bias [38]. This experimental heterogeneity required critical data processing to ensure statistically meaningful assumptions to drive biological interpretation and compile gene signatures. For this, we used a standardized workflow to reduce as much as possible the technical variations between datasets. Specifically, this workflow applied i) the same normalization method for the experiments having the same microarray chips, ii) the same quality control criteria to discard poor-quality samples, iii) the same aggregation method for summarizing probesets into single genes and iv) the same filtering in all datasets. The filtering of the datasets was based on the same significance criteria which included: a minimum number of differentially expressed genes, the presence of genes known to be differentially expressed between quiescent and activated states from previous studies and a similarity measure among the datasets. Table 1 summarizes the main biological and experimental variations in this study, as well as the technical differences present in the datasets.

Three new sets of microarrays of quiescent versus activated satellite cell are reported here (see Table 1). The first one is part of a developmental and postnatal series that was reported previously [16] (E12.5 vs. E17.5), and here P8 (postnatal day 8, *in vivo* proliferating) and 4-5 week old (quiescent) mice were compared. The second one is based on previously reported differences in quiescent and proliferating cell states in subpopulations of MuSCs (*Quiescent*: dormant, top 10% GFP+ cells vs. primed, bottom 10% GFP+ cells isolated from *Tg:Pax7-nGFP* mice; *Proliferating*: 3 days post-injury [3]). The third dataset is based on previous observations that the *Notch* intracellular domain (NICD) when expressed constitutively (*Myf5 ^Cre^: R26 ^stop-NICD^*) in prenatal muscle progenitors leads to cell-autonomous expansion of the myogenic progenitor population (*Pax7*+/*Myod*-) and the absence of differentiation, followed by premature quiescence at late foetal stages (E17.5) [16]. Here, E17.5 (quiescent) and E14.5 (proliferating) prenatal progenitors were compared. Except for our datasets *Quiescent(adult)/Activated(P8)* and *Foetal_NICD[E17.5/E14.5],* all the studies were conducted on adult mice (male and female) with ages ranging from 8 weeks to 6 months.

While all datasets shared similar cell states (quiescent (QSC) and activated (ASC) satellite cells), the experimental procedures varied between studies. Activation of cells, for instance, was achieved in different ways: i) *in vitro,* by culturing freshly isolated MuSCs for several days, ii) *in vivo*, by extracting ASCs from an injured muscle. Furthermore, for *in vivo* activation, several techniques were used to induce the injuries: BaCl_2_, or the snake venoms cardiotoxin or notexin. Cell extraction protocols also varied among the different studies: i) using transgenic mice expressing a reporter gene that marks satellite cells (several alleles) or, ii) using a combination of antibodies targeting surface cell antigens specific to satellite cells (several combinations, see Table 1). Finally, the nine datasets examined in this study date from 2007 to 2016. During this period, microarray technologies evolved and the different chips available may introduce yet another source of variation among the compared datasets. To carry out a statistically meaningful analysis of these extensively heterogeneous datasets, critical data processing was required to interpret gene signatures as described in the workflow (Fig. 1).

### The number of differentially expressed genes varies significantly among different datasets

A total of 32 samples from ASCs and 34 samples from QSCs from the nine datasets were analysed. After the quality control, one sample from the GSE38870 dataset was considered to be an outlier and was not included in the final analysis.

The number of significantly up and down regulated genes (DEGs) resulting from the differential expression analysis of the quiescent with respect to the activated states were noted (Additional file 2: Table S1). DEGs were identified as having |logFC| < = 1 and a false discovery rate FDR = = 0.05. The statistical analysis was performed at the probeset level, and only those probesets matching to genes are reported. On average, the datasets exhibited 1548 up-regulated genes with a standard deviation of 1173 genes. The number of down-regulated genes corresponded to 2122, with a standard deviation of 1658 genes. The lowest number of DEGs belonged to the *Foetal_NICD[E17.5/E14.5]* dataset (39 up, 136 down), while the highest number of DEGs belonged to the GSE70376 dataset (4367 up, 6346 down). Additionally, an analysis of the distribution of the logFC across the datasets revealed that there were significant differences among the ranges and shapes of such distributions for each dataset (Additional file 3: Fig. S2).

### Gene set level analysis reveals common underlying biological processes across the datasets

Despite the great difference among the number of DEGs for the different sets, clear trends among the significantly enriched pathways were found (Fig. 3a). This heatmap shows each dataset as a column and each enriched gene set as a row. The gene set collection that was tested for enrichment corresponds to the Hallmark gene set collection from MSigDB [39]. Enriched gene sets corresponding to over-expressed genes are shown in red, while enriched gene sets that were generally abundant in under-expressed genes are shown in blue. Out of the 11 datasets, GSE38870 stood as an outlier for both over and under-represented gene sets compared to the rest. For the other 10 datasets, most of them showed an enrichment in the quiescent state for the TNFA_SIGNALING_VIA_NFKB pathway (9 datasets), while 8 datasets were enriched in UV_RESPONSE_DN, IL6_JAK_STAT3_SIGNALING, APICAL_SURFACE and KRAS-SIGNALING_DN pathways. Similarly, the 10 datasets shared similar trends for under-expressed genes in the pathways MYC_TARGETS_V1, E2F_TARGETS, G2M_CHECKPOINT, and OXYDATIVE_PHOSPORYLATION, all of which are expected to be absent in the quiescent state. In total, two subnetworks corresponding to 8 under- and 15 over-expressed enriched gene sets could be distinguished (Fig. 3b). A network representation of the top 3 most commonly found enriched gene sets (nodes, thick outlined circles) is shown in Fig. 3b for the over-expressed (TNFA_SIGNALING_VIA_NFKB, UV_RESPONSE_DN, IL6_JAK_STAT3_SIGNALING) and under-expressed (MYC_TARGETS_V1, E2F_TARGETS, G2M_CHECKPOINT) categories. The size of each node corresponds to the total number of times that gene set was enriched in all the datasets, and the thickness of the interconnecting lines is proportional to the number of genes shared between connected nodes. Gene sets sharing less than 10% of their genes are not shown. We noted also that different gene sets had a varying number of genes in common (Fig. 3b); if the gene overlap were large, those gene sets (and their corresponding biological functions) will likely be also affected (i.e. activated or repressed). For the 3 most common enriched gene sets with under-expressed genes, for example, we noted that gene set MYC_TARGETS_V1 shares most of its genes with gene sets E2F_TARGETS and G2M_CHECKPOINT. This suggests that the three categories represented by these gene sets had an interplay of genes that displays them as all under-expressed.

**Fig. 3.**
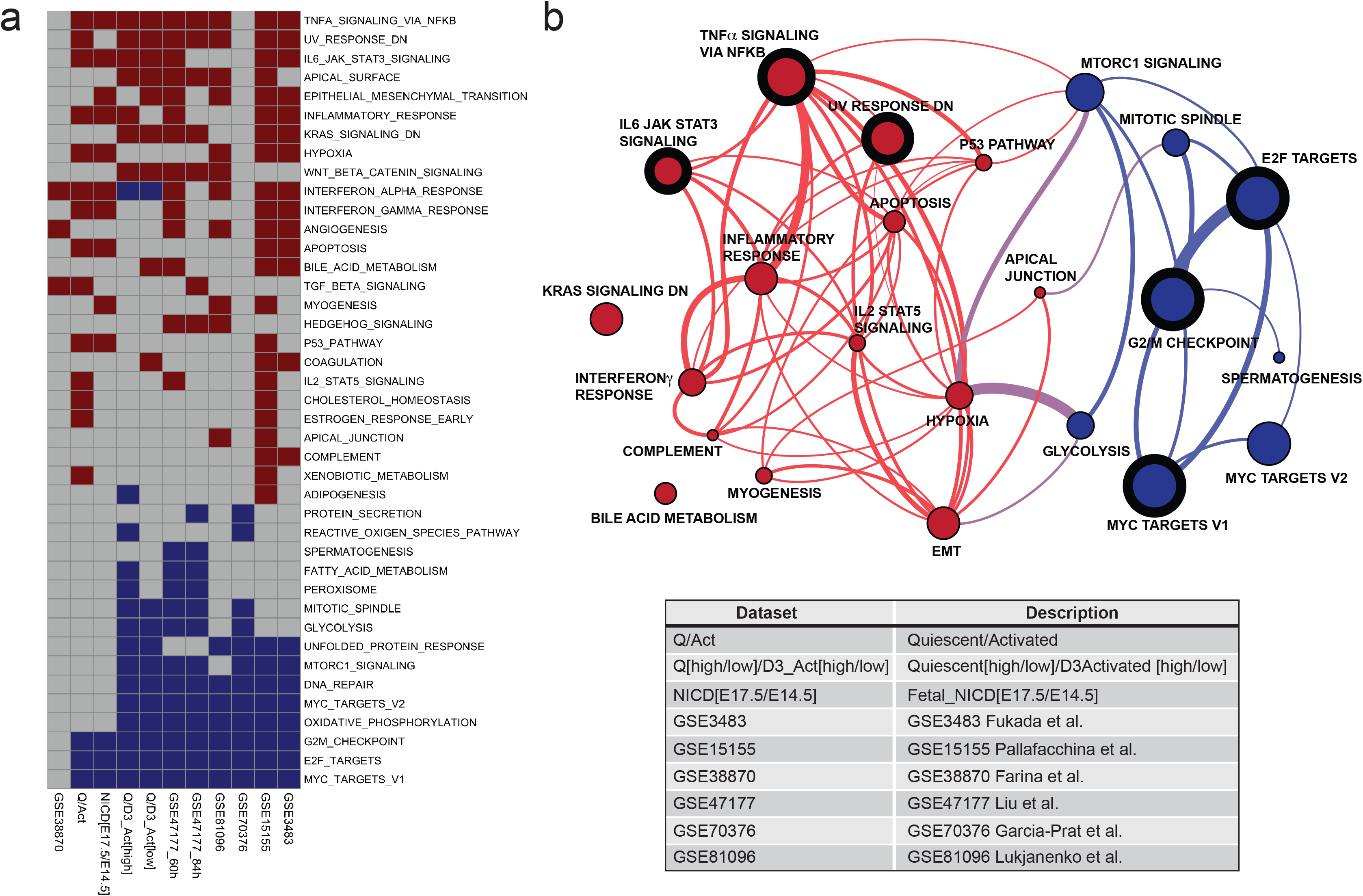
Enriched gene sets across individual datasets. Enriched (over-represented) gene sets with over-expressed genes are shown in red; enriched gene sets with under-expressed genes are shown in blue. a) Gene set enrichment profiles using the Hallmark gene set collection from MSigDB [39], each row corresponds to a gene set, and each column corresponds to a dataset. b) Network representation of 3 most common over and under-expressed gene sets (denoted by thick border on node) along with gene sets sharing genes with them (connector lines). Nodes represent gene sets with a node circle size proportional to the number of times the gene sets appear as enriched in the different datasets (see Fig. 3a). Thickness of the connecting lines is proportional to the number of shared genes between nodes.

### Determining a quiescent transcriptional signature among all datasets

To determine a consensus quiescent signature from the datasets we compared the genes found to be differentially expressed within each dataset, in order to identify genes commonly up- or down-regulated in the quiescent state. Although the aforementioned technical and experimental heterogeneity could introduce noise in this analysis, such variation was distinguishable from the more stable, underlying common quiescent signature. Given that the distribution and ranges of the logFCs varied so drastically between datasets (Fig. S2), a single FC (fold change) threshold could not be chosen to be used for all datasets. Thus, for the combinatorial analysis approach, we set out to maximize the number of differentially expressed genes common to all the datasets that were considered, where only the adjusted p-value was used as a threshold to define DEGs. However, even in this low constrained scenario, combining all the datasets together resulted in very few overlapping genes: 12 up (*Arntl*, *Atf3*, *Atp1a2*, *Cdh13*, *Dnajb1*, *Enpp2, Ier2*, *Jun*, *Nfkbiz*, *Rgs4*, *Usp2*, *Zfp36*) and 1 down (*Igfbp2*). Alternatively, when certain datasets were excluded from the analysis, the number of DEGs increased (Fig. 4a).

**Fig. 4.**
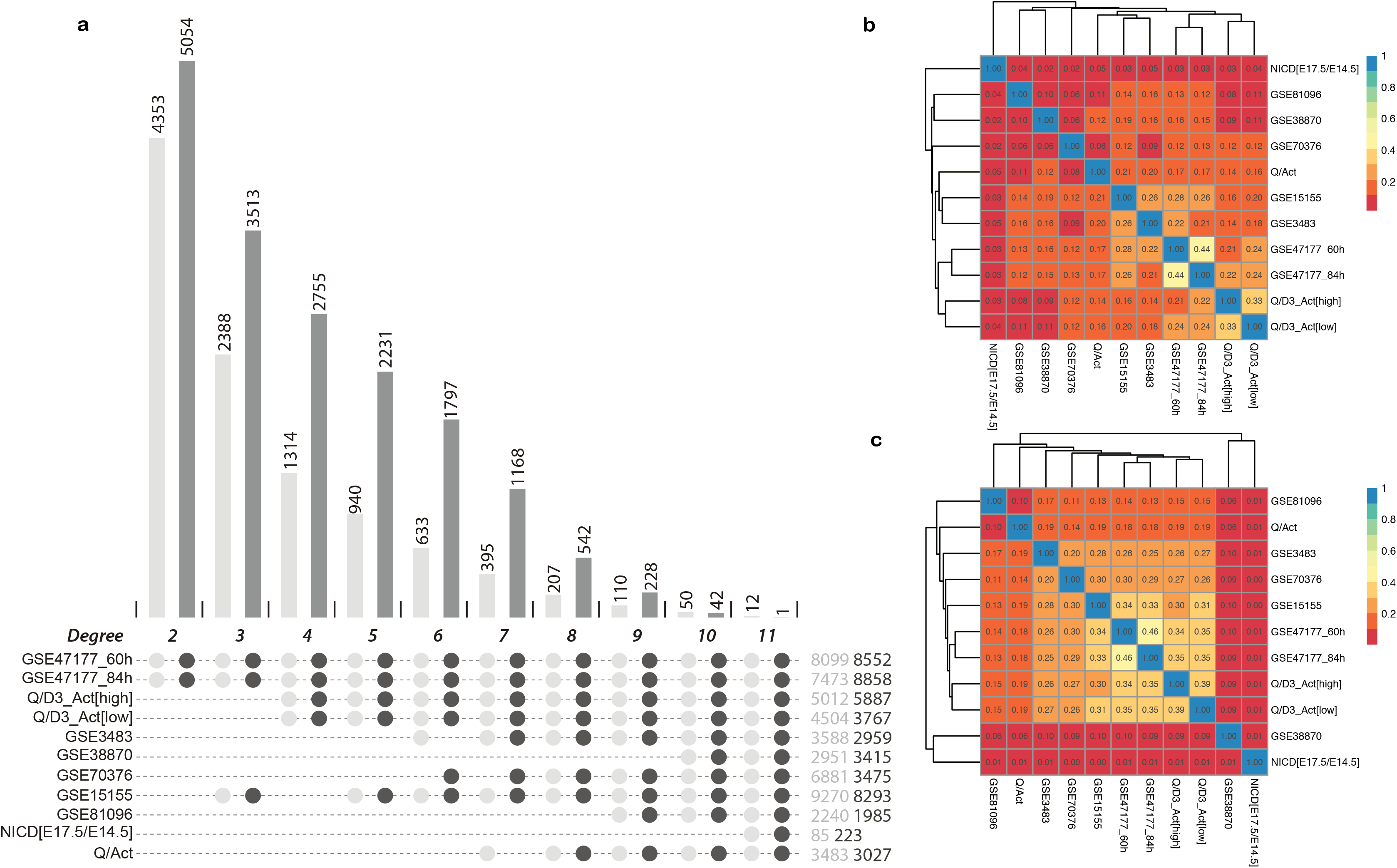
Different combinatorial landscapes result in different degrees of stringency for the list of genes defining the quiescent state of MuSCs. a) Barplot indicating the number of overlapping differentially expressed genes (DEGs) for each best combination of intersections, from degree 2 to 11. The dots underneath the barplot indicate the datasets included in the intersections. The total number of up (UP) and down (DOWN) DEGs for each dataset are indicated in light grey and dark grey, respectively. b) and c) are colored matrices showing the Jaccard index between each pair of datasets, for UP DEGs and DOWN DEGs, respectively. Dendrograms show the hierarchical clustering using the Jaccard index as euclidean distance.

### Combinatorial assessment of datasets according to significance and similarity criteria

To find the best combination of datasets defining a consistent and sufficiently large quiescent signature, we ranked them according to their significance. This significance was determined according to an ensemble of criteria. First, the dataset should have a minimum number of DEGs. Our *Foetal_NICD[E17.5/E14.5]* dataset, for instance, had only 250 DEGs (Table S1), and using it in the analysis resulted in a dramatically low number of overlapping DEGs (Fig. S3). A second criterion was the presence of genes known to be differentially expressed between quiescent and activated states from previous studies. In this case, datasets GSE38870 and GSE81096 did not meet this criterion, since they lacked genes known to be associated with or regulating the quiescent state such as *Calcr, Notch1*, *Chrdl2, Lama3, Pax7 and Bmp6* genes (unpublished data, see Fig. 7; [51][63][64]). As a third criterion, we used the dataset similarity, which was assessed using the Jaccard Index (JI), and a matrix of the JIs for the up- and down-regulated genes was generated (Figs. 4b, c, respectively). In both matrices, the closest pairs of datasets were GSE47177 at 60 hours and GSE47177 at 84 hours (JI = 0.46 and 0.44 for the up and down regulated genes, respectively), followed by the second pair of closest sets Quiescent [high] / D3Activated [high] and Quiescent [low] / D3Activated [low] (JI = 0.39 and 0.33, for up and down regulated genes, respectively). The observation that the first two closest datasets belonged to studies originating from the same laboratory underscores the impact of technical biases. The hierarchical clustering of the Euclidean distance of the Jaccard indexes shows that for up and down regulated genes, the datasets *Foetal_NICD[E17.5/E14.5]*, GSE38870 and GSE81096 had a tendency to not group with the rest of the datasets. In addition to these criteria, others can be used to assess the significance of the datasets. Choosing the datasets according to the activation or extraction method of the cells, for example, would result in a more stringent ensemble of datasets.

Taking into account the dataset significance (based on number of DEGs and presence of some reported quiescent markers) and the low extent of overlap between *Foetal_NICD[E17.5/E14.5]*, GSE38870 and GSE81096 datasets with respect to the remaining datasets, these three datasets were excluded from the multi-dataset analyses. The final ensemble comprised the eight remaining datasets which had 207 and 542 genes commonly up- and down-regulated, respectively (Fig. 4a). To further characterise these commonly regulated genes, we performed an over-representation analysis (ORA) of the gene sets. An enrichment was detected for the 207 commonly up-regulated genes in seven different Hallmark gene sets (Fig. 5a). Some genes were shared among different pathways (e.g. *Atf3* and *Il6* were found in six different gene sets), while others were found in one gene set only (e.g. *Tgfbr3*, *Spsb1*). These results are consistent with the individual gene set enrichment analysis (see Fig. 3) emphasizing that these genes reflect the global traits associated with the quiescent state. Note that only a fraction of the 207 genes was found in known existing gene sets (57/207), leaving about three quarters of the commonly up-regulated genes not associated with any existing gene set. This finding was expected given that a quiescent signature is yet to be defined, and thus current gene sets lack such annotations. To facilitate the analysis of transcriptomes as described here, we have developed an online interactive tool called *Sherpa* (Fig. 6). *Sherpa* allows users to perform analyses on individual and on multiple datasets. Each individual dataset analysis involves the identification of differentially expressed genes, comparison of the expression of selected genes in the quiescent and activated states through tables, heatmaps and volcano plots, exploration of the distribution of the samples according to their variability through Principal Component Analysis, and cluster analysis. The multiple dataset analysis allows the comparison of selected datasets according to the commonly differentially expressed genes. All of these analyses are interactive, as they allow the user to select the thresholds of fold change (logFC) and false discovery rate (adj. P-value).

**Fig. 5.**
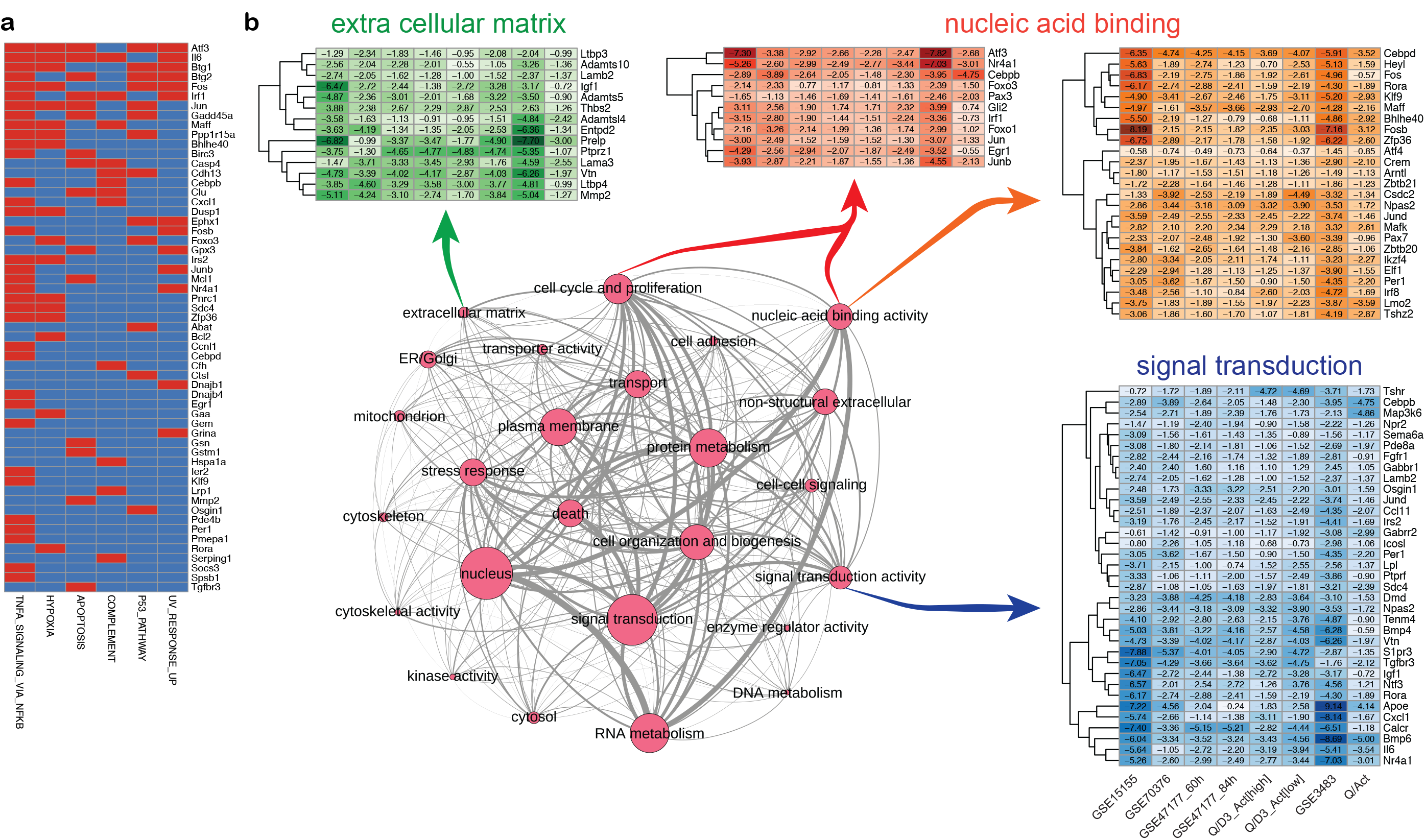
Gene expression of differentially expressed genes (DEGs) in MuSCs. a) Binary heatmap of the over representation analysis. Each column represents one enriched (over-represented) gene set, and each row corresponds to a gene. Red cells indicate the presence of the corresponding gene in a given gene set. b) Network representation of 39 GOSlim terms used to characterize the commonly regulated genes in MuSCs. Nodes represent gene sets with a node size proportional to the gene set size. Edges indicate that genes are shared among the gene sets. Thickness of the edge is proportional to the number of shared genes. Also shown are the heatmaps of logFC for genes belonging to extracellular matrix, nucleic acid binding and cell cycle and proliferation, nucleic acid binding and signal transduction activity, respectively. Each row corresponds to a gene and each column corresponds to a dataset. Dendrograms show hierarchical clustering using the euclidean distance.

**Fig. 6.**
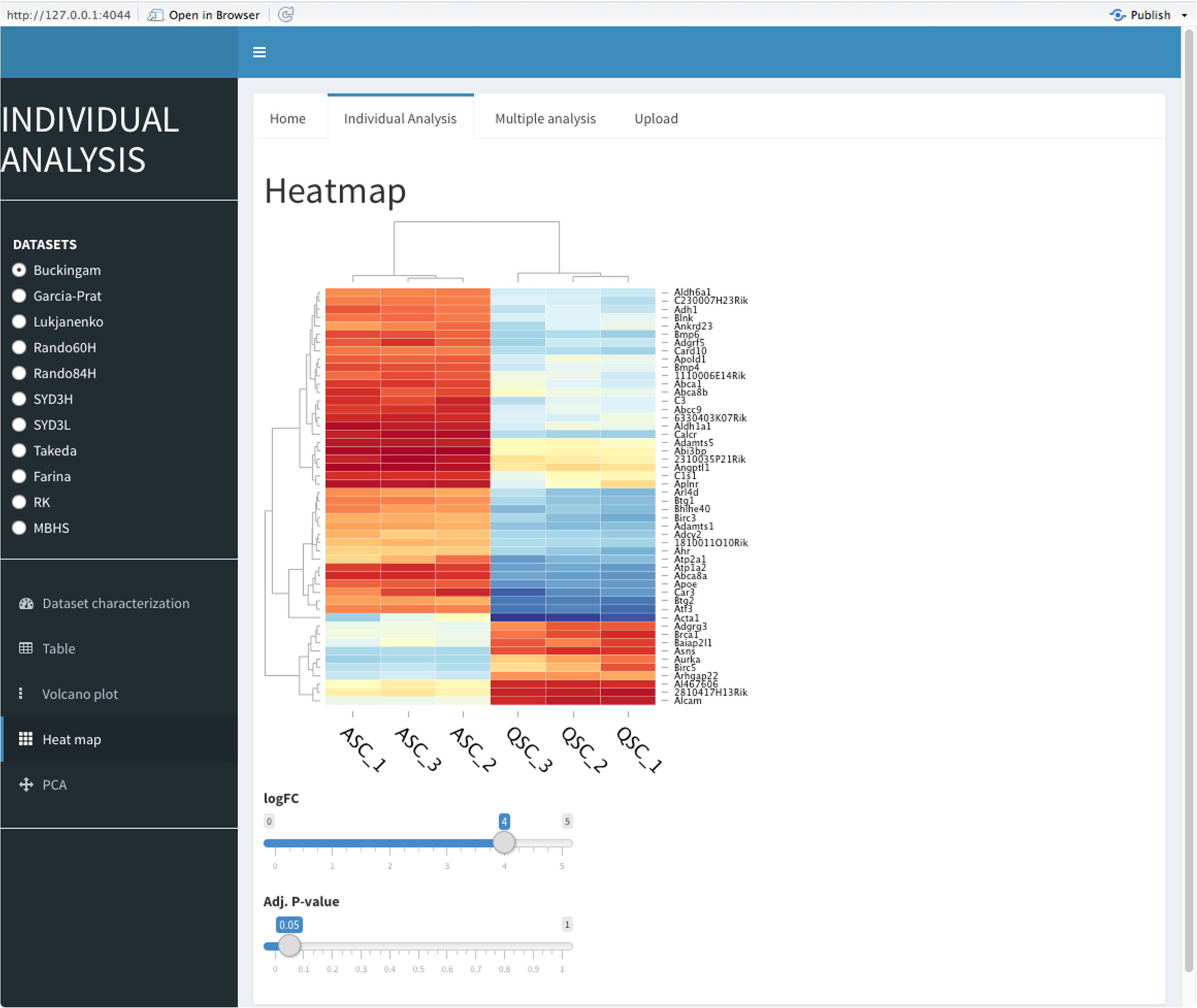
Snapshot of the interactive web application for transcriptomic data exploration and comparison. *Sherpa* (http://sherpa.pasteur.fr) allows users to perform individual dataset and multiple dataset analysis. In the individual dataset analysis (shown), the user chooses the dataset for which the analysis is to be performed. Then, it is possible to identify differentially expressed genes (e.g. Volcano plot), compare the expression of selected genes in the quiescent and activated state (e.g heatmap, as shown in Figure), the distribution of the samples according to their variability (Principal Component Analysis). All these analyses are interactive, as they allow the user to set the thresholds of fold change (logFC) and false discovery rate (adj. P-value).

To assign a global function to the commonly regulated genes, we annotated them using GOSlim terms, which summarize broad terms based on Gene Ontology (GO) terms [40]. To identify categories of genes, heatmaps of the logFC in the different datasets for a subset of the 207 UP genes belonging to extracellular matrix, nucleic acid binding activity (+/- cell cycle proliferation) and signal transduction activity were generated (Fig. 5b). Unexpectedly, genes associated with cell cycle proliferation, such as *c-Fos*, *c-Jun*, were up-regulated in the quiescent cell analyses in all datasets (Fig.7a). To verify the transcriptional relevance of these genes in quiescent cells, we used a protocol to isolate MuSCs in which a short fixation (PFA) treatment was performed prior to harvesting the cells to arrest *de novo* transcription during the isolation protocol (see Methods). Then, expression level quantification was assessed at the transcript (RT-qPCR) and protein (Western blot) level at different time points after isolation for a number of genes (Fig. 7b, c). Notably, quantifications of *c-Jun, Jun B* and *Jun D* levels showed that at time 0 (+PFA), these genes were not detected in quiescent cells, neither at the mRNA (right panel), nor at the protein (left panel) level (Fig. 7c). However, these genes were up-regulated using conventional MuSC isolation protocols that take several hours. As a control, PFA treatment after cell isolation had no effect on this expression pattern (Fig. 4S). This rapid up-regulation was then followed by a decline in expression levels of these genes (Fig. 7b, c), suggesting that this is the result of a stress response that is associated with the isolation procedure.

**Fig. 7.**
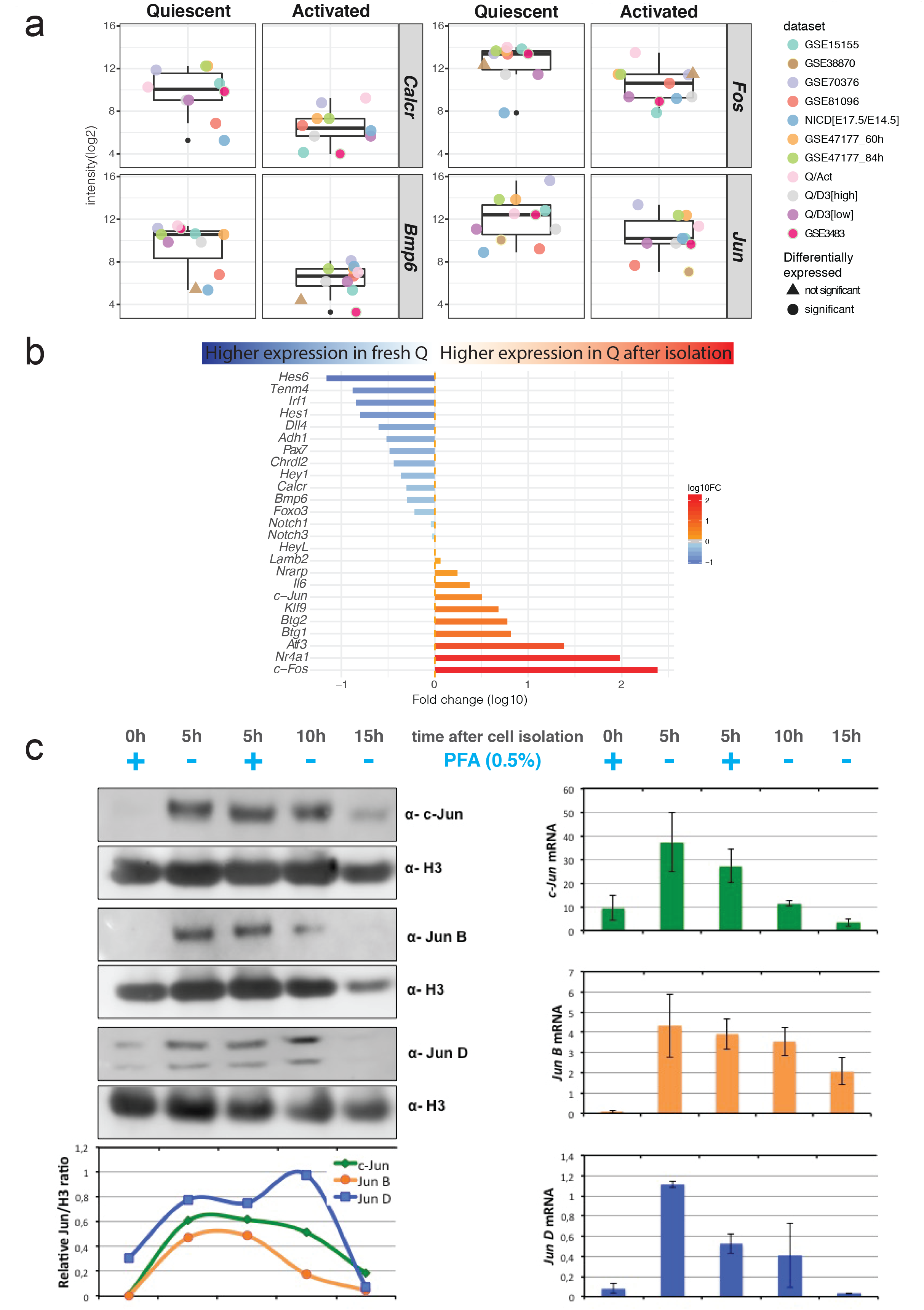
Direct comparison of fixed and unfixed MuSCs identify immediate response genes not present the in vivo state. a) boxplots of four examples of genes found commonly upregulated in QSCs in the different datasets showing the distribution of intensities values in QSCs and ASCs. Colored dots indicate each dataset. Shape of the dot indicates whether the gene is significantly differentially expressed or not. b) Fold change of mRNA (log10) between 0h+PFA and 5h+PFA. Blue bars indicate a higher expression in 0h+PFA condition, the red bars indicate a higher expression in 5h+PFA condition. Color intensities are proportional to the fold change. c) *c-Jun*, *Jun B* and *Jun D* protein levels from MuSCs at 0, 5, 10, 15h after isolation (with and without PFA treatment) were measured by Western blotting and band intensities were quantified by densitometric analysis with the ImageLab software (right). Basal levels of c-Jun, Jun B and Jun D mRNA from MuSCs at 0, 5, 10, 15h after isolation (with and without PFA treatment) were measured by real-time PCR (left).

## Discussion

The transcriptome analysis and pipeline, as well as the *Sherpa* interface that we describe here, allow multiscale comparisons across divergent datasets that are heterogeneous in platform and biological condition. Notably, this pipeline allowed the examination of 11 datasets, including 3 novel transcriptomes from our work, as well as the identification of a variety of gene sets that appear in common with the majority of the datasets. To perform this analysis, it was necessary to standardize as much as possible every step of the analysis to attenuate the impact of heterogeneity inherent in all of the datasets due to experimental, biological and technical variations. These varying conditions lead us to perform a combinatorial assessment of the individual datasets according to their significance and similarity criteria.

Variations in datasets are not unique to the study of muscle stem cells. Indeed, the last decades have witnessed many efforts to analyse microarray data to provide relevant gene signatures. In cancer biology, for example, gene markers were sought either for prognosis, i.e. lists of genes able to predict clinical outcome [41] or for molecular subtyping, i.e. list of genes able to classify different subtypes of a disease [42][43]. However, even if markers performed well, gene signatures derived from studies on the same treatments and diseases often resulted in gene lists with little overlap [44]. In other cases, the signatures proved to be unstable, having other gene lists on the same dataset with the same predictive power [45]. These observations suggest that such signatures may include causally related genes, i.e. downstream of the phenotype causing genes and that these gene lists may share the same biological pathways [46].

Gene Set Enrichment Analysis (GSEA) has become an efficient complementary approach for analysing *omic* data in general and GEPs in particular [47][46][48]. It shifts the expression analysis from a *gene* space to a *gene set* space, where genes are organized into gene sets according to a common feature, such as a functional annotation (e.g. a Gene Ontology term) or a specific metabolic pathway (e.g. a KEGG pathway). In this way, it incorporates previously existing biological knowledge to drive and increase interpretation, while offering greater robustness and sensitivity than gene level strategies [46][49][50].

In spite of the heterogeneity in datasets examining quiescent muscle stem cells, we were able to identify genes that were consistently up- and down-regulated among the different datasets. The final multiset analysis comprised eight datasets which had 207 and 542 genes that were commonly up- and down-regulated, respectively. Moreover, the gene set enrichment analysis of the individual datasets showed striking similarities on the over- and under-represented gene sets. These gene sets, which summarize and represent well-defined biological states and processes in the cells, were shared among the different datasets. They include an over-representation of genes in the TNFa pathway via NFKb signaling, Il6-Jak-Stat3 signaling, and the apical surface processes, and an under-representation of MYC and E2F targets, and genes associated with the G2M checkpoint and oxidative phosphorylation. Some markers such as *Calcitonin receptor (Calcr)*, *Teneurin4* (*Tenm4*), and stress pathways identified previously were also present in our analysis [51][52][11]. However, we also report that virtually all datasets contained genes that would be expected to be present during activation or cell cycle entry, such as members of the *Fos* and *Jun* family previously identified as immediate early stress response genes [53]. Using a novel isolation protocol (Mashado et al., in press; P. Mourikis, F. Rélaix, personal communication) based on the notion that tissues that are fixed prior to processing result in stabilized mRNA [18], we validated the expression of several genes including *Calcr* and *Teneurin4* (*Tenm4*) as true quiescent markers. In contrast, we show that *Fos* and *Jun* transcripts, and *Jun* family proteins are not present at significant levels *in vivo*, but are robustly induced within 5 hours, the average processing time taken for isolation by FACS of MuSCs. These results are concordant with a recently published paper in which immediate early and heat-shock genes were rapidly up-regulated during the cell isolation procedure [62]. We propose that these and other stress response genes mitigate the quiescent to activation transition that accompany the initial steps of exit from G0.

Given these unexpected findings, the comparison of transcriptomes of MuSCs from a fixed/*in vivo* state with those that were described here would be important to delineate homeostatic vs. immediate early response genes. For that purpose, *Sherpa* allows the integration of datasets from fixed samples, or other methodologies, when they will be available. Beyond the present findings, we propose that all transcriptome data obtained from cells isolated from solid tissues, which require extensive enzymatic digestion and processing before isolation of RNA, need to be re-evaluated to distinguish those genes that are induced by the isolation procedure.

In addition to generating this open access compendium of GEPs, we provide a standardized pipeline that sets the basis for a multi-set analysis for an effective and systematic comparison of individual datasets. Analysing multiple datasets provides generalized information across different studies [37][54]. The cancer field was a pioneer in combining several works [55] [56] and other fields, such as neurodegenerative diseases [57][58] and regulatory genomics have successfully adopted this strategy [59]. The multidimensional approach presented here offers increased power, due to the higher sample size and increased robustness, by highlighting variations in individual studies results [36][60]. Such variations are the consequence of the high level of noise and artefacts, and are typically associated with microarray data [61].

## Conclusions

Here we compile the first comprehensive catalogue of gene expression data of myogenic cells across distinct states and conditions, providing a global perspective on quiescence. An extensive comparison of the transcriptomic profiles of mouse skeletal muscle stem cells in quiescent and activated states resulting from nine datasets revealed common features among the different studies from other features which are more specific to the individual datasets. In spite of heterogeneities across platforms, we were able to identify genes that were consistently up and down regulated among the different datasets. By doing so, we developed and made available an open-access interactive exploratory tool called Sherpa (SHiny ExploRation tool for transcriPtomic Analysis) that allows statistically valid analyses and systematic comparisons that cannot be performed directly on the datasets. Finally, by obtaining mRNA directly from fixed muscle tissue for empirical testing of genes present during quiescence *in vivo*, we identified immediate early expressed stress response genes that were present in all datasets due to the isolation and processing protocols used previously for solid tissues.

## List of abbreviations

ASCs: Activated Satellite Cells
DEGs: statistically Differentially Expressed Genes
FACS: Fluorescence Activated Cell Sorting
FC: Fold Change
FDR: False Discovery Rate
FSC: Functional Scoring Method
GEO: Gene Expression Omnibus
GEPs: Gene expression Profiles
GFP: Green Fluorescent Protein
GO: Gene Ontology
GSEA: Gene Set Enrichment Analysis
KEGG: Kyoto Encyclopedia of Genes and Genomes
logFC: logarithm of the Fold Change
MuSC: skeletal muscle stem (satellite) cells
NICD: Notch Intracellular Domain
NUSE: Normalised Unscaled Standard Errors
ORA: Over Representation Analysis
P8: postnatal day 8
PFA: paraformaldehyde
QSCs: Quiescent Satellite Cells
RLE: Relative Log Expression
RMA: Robust Multi-Array Average expression measure
Sherpa: SHiny ExploRation tool for transcriPtomic Analysis
TA: Tibialis Anterior
TCGA: The Cancer Genome Atlas

## Declarations

### Ethical Approval and Consent to Participate

Not applicable.

### Consent for publication

Not applicable.

### Availability of supporting data

The generated transcriptome datasets are available from the corresponding author on reasonable request. Public datasets are available at https://www.ncbi.nlm.nih.gov/geo/ under their corresponding identification number.

### Authors' contributions

NP, SM, ST analysed and interpreted the data regarding the gene expression profiles. SY, MBB, HS and RS performed experiments to generate transcriptome data. FP contributed to data analysis. DDG and FP performed validation experiments of candidate genes. All authors read and approved the final manuscript.

## Acknowledgements

We would like to thank K. Soni and U. Borello for their assistance in the early stages of this work, and P. Mourikis and F. Relaix for communicating unpublished results and sharing protocols. We also acknowledge the Flow Cytometry Platform of the Technology Core-Center for Translational Science (CRT) at Institut Pasteur for support in conducting this study and the microarray platform at Institut Cochin.

## Supplementary Table and Figure Legends

**Fig. S1. Quality controls and data sample distribution for Quiescent [high/low] / D3Activated [high/low] dataset.** a) Relative Log Expression (RLE) and b) Normalised Unscaled Standard Errors (NUSE) plots for the D3P7 dataset show that as expected for good quality data, RLE median values are centered around 0.0 while the median standard error should be 1 for most genes in the NUSE plots. Sample distribution is distributed according to status (D3H: activated, high; D3L: activated, low; QH: quiescent, high; QL: quiescent, low) using c) Principal Component Analysis and d) hierarchical clustering of the Euclidean distance.

**Fig. S2. Violin plots of the logFC distribution for each individual dataset.** Density plots of the logFC (|logFC| = 1 in red; |logFC|< 1 in blue.

**Fig. S3. Effect of adding NICD[E17.5/E14.5] dataset on the best combinations of datasets.** Best combination of datasets was determined by the bigger overlap between n ( = degree) datasets. n varies from 1 (the dataset containing the most DEGs) to 10 (all the datasets except NICD[E17.5/E14.5]). Blue bars/numbers indicate the number of best overlap between the n datasets, orange bars/numbers indicate, the extent of overlap when NICD[E17.5/E14.5] was added to this best combination.

**Fig. S4. Effect of PFA treatment at different time points in the experimental procedure.** a) Schematic showing the simplified experimental procedure of cell fixation before or after muscle dissociation/cell sorting. “0h” refers to cells fixed prior to muscle dissociation and cell sorting, while “5h” refers to cells that undergo muscle dissociation and cell sorting prior any treatment (+/- PFA). b) Barplot showing the effect of PFA treatment after muscle dissociation/cell sorting. Bar represent the fold change (in Log10) of expression between freshly isolated QSCs (0h + PFA) and QSCs after muscle dissociation/cell sorting with or without PFA treatment, 5h +PFA and 5h –PFA, respectively.

**Table S1. Identified differentially expressed genes in the quiescent satellite cell condition for the 9 datasets.**

**Table S2. Primers used for validation of gene expression by RT-qPCR.**

